# Identification and assessment of the antibacterial activity of *Vigna mungo* and Rhizobacteria

**DOI:** 10.1101/2021.10.23.465580

**Authors:** Bharat Kwatra

## Abstract

ZnO nanoparticles have received a lot of interest in recent years due to their unusual features. Antimicrobial properties of ZnO NPs However, the qualities of nanoparticles are determined by their size and form, making them suitable for a variety of applications. The current work looks at the synthesis, characterization, and antibacterial activity of ZnO NPs produced by Vigna Mungo and Rhizobacteria. Rhizobacteria isolated from V. mungo root nodule were morphologically, biochemically, and molecularly examined and identified as Rhizobium sp. strain and Bacillus flexus strain. The GC-MS analysis of methanol leaf extract of V. mungo was performed to detect and identify bioactive chemicals, and this indicated phytol as an antibacterial agent, while Squalene and Alpha – tocopherol had antioxidant and anti-tumour properties. Agar well diffusion experiment was used to determine the antibacterial properties of ZnO nanoparticles and Vigna Mungo leaf extract. This approach is widely documented, and standard zones of inhibition for sensitive and resistant values have been defined. The results demonstrated that both methanol extract and zinc oxide nanoparticles have strong antibacterial efficacy against the majority of the pathogens examined.

## INTRODUCTION

Nanotechnology is one of the most active fields of research in modern materials science. Based on certain qualities such as size, distribution, and shape, nanoparticles display new or better properties. In recent years, there have been tremendous advances in the science of nanotechnology, with several approaches created to synthesis nanoparticles of specified form and size based on specific requirements. The number of new uses for nanoparticles and nanomaterials is continually expanding [1]. Metal oxides include titanium dioxide (TiO2), indium (III) oxide (In2O3), zinc oxide (ZnO), tin (IV), and silicon dioxide (SiO2), with ZnO being the third most widely produced metal oxide after SiO2 and TiO2. ZnO is an inorganic substance with unique qualities such as semiconductor, a broad spectrum of radiation absorption, piezoelectricity, pyroelectricity, and high catalytic activity. Furthermore, due to its non-toxic qualities, ZnO has been designated as “Generally Recognized as Safe” (GRAS) by the US Food and Drug Administration (FDA 21CFR182.8991). As a result, ZnO NPs are now widely used in electronics, optics, biomedicine, and agriculture. Nature has evolved several techniques for the synthesis of nano and micro-length scaled inorganic materials, which have contributed to the establishment of a relatively new and mostly unknown area of research centred on nanomaterial biosynthesis. The use of bio organisms in synthesis is consistent with green chemistry concepts. Because of their low water requirements and capacity to endure environmental stress, pulse crops have a prominent position in the farming system. Through symbiosis with Rhizobium, pulses may repair atmospheric nitrogen. Pulses are renowned as soil fertility restorers due to their unique capacity to mobilise insoluble soil nutrients through the deep root system and bring about qualitative changes in soil physical properties [2, 3]. The plant’s flavonoid inducers play an important part in the process by prompting the Rhizobium to generate certain Nod factors that activate the host symbiotic mechanisms required for root hair infection and nodule growth.The antibacterial activity of ZnO nanoparticles is determined by surface area and concentration, with no impact from crystalline structure or particle shape. The smaller the size of ZnO particles, the greater their antibacterial activity. Thus, the greater the concentration and surface area of the nanoparticles, the greater their antibacterial activity. The mechanism of antibacterial action of ZnO particles is currently unknown. Some researchers argued in their study that the production of hydrogen peroxide is the primary cause of antibacterial activity, while others suggested that particle attachment on the bacterium surface is caused by electrostatic forces could be another factor [5, 6].

## 2. MATERIALS AND METHODS

### 3.1. Preparation of plant extract

Fresh leaves of *Vigna Mungo* were collected from the local regions of Thiruvananthapuram district, Kerala and washed several times with sterile distilled water and then dried using a hot air oven for 2- 3 days and grinded to form powder. The powdered plant sample was sequentially extracted using three solvents namely hexane, butanol and methanol. The extracts were dried and the solvent-free extracts were stored in an air tight container for further study.

#### 3.1.1. Isolation of *Rhizobacteria* from *Vigna Mungo*

Healthy unbroken pink nodules were selected for the isolation. The Nodules were picked with sterile forceps. The collected nodules were kept in sterile polythene bags and transported to the laboratory for further investigation ^[7]^. They were dipped in 0.1 % HgCl_2_ or 3-5 % H_2_O_2_ for five minutes for surface sterilization of nodules. The sterilized root nodules were crushed by adding 1 mL of sterile distilled water. This suspension was serially diluted up to 10^−5^. The diluted suspensions 10^−2^-10^−4^ were selected, and 0.1 mL of suspension was inoculated in petri plates containing sterile Yeast Extract Mannitol Agar medium (YEMA) with congored. The inoculated plates were incubated at 30 **±** 2°C for four days.

#### 3.1.2. Morphological characterization of *Rhizobacteria*

The bacterial isolates were grown on YEMA medium. After 24 to 72 hours, the colony morphology of isolates was studied with special consideration to the color, size of colonies, and their margins.

#### 3.1.3. Biochemical characterization of *Rhizobacteria*

The various biochemical characteristics viz., gram staining, motility test, indole production test, MR test, VP test, citrate utilization test, Oxidase test, GPA test, lactose assay, keto lactose test, starch hydrolysis test, gelatin hydrolysis test, fluorescent assay, TSI test, Urease test, Catalase test and nitrate reduction test were carried out ^[8-11]^.

#### 3.1.4. Molecular characterization of *Rhizobacteria*

The identification of virus requires metagenomic sequencing (the direct sequencing of the total DNA extracted from a microbial community) due to their lack of the phylogenetic marker gene 16S.

### 3.2. Antibacterial activity

The isolated three bacterial strains such as *Bacillus velezensis, Bacillus zanthoxyli* and *Klebsiella ozaenae* were used for determining the antibacterial activity of different plant extracts and nanoparticles.

### 3.3. Minimum inhibitory concentration (MIC)

100 µL of different concentrations (0.02µg/mL, 0.04µg/mL, 0.06µg/mL and 0.08µg/mL) of nanoparticles were added to the wells and allowed to diffuse at room temperature for 2 hours. The cultures were incubated at 37°C for 24 hours ^[13]^.

### 3.4. Minimum bactericidal concentration (MBC)

The nanoparticles were diluted into various concentrations (0.06, 0.07, 0.08, 0.09, 0.1 and 0.5µg/mL) in sterile nutrient broth (5 mL) in test tubes. The 20 µL of *Bacillus velezensis* and *Klebsiella ozaenae* were added to all tubes and incubated at 37°C for 24 hours. The lowest concentration in which has no single colony bacterial growth was taken as MBC ^[14]^.

### 3.5. GC-MS analysis

GC-MS analysis was carried out on a resolution GC-Agilent 7890 A and MS- Agilent 5975 C. The column used was DB5- MS (30m x 25mm x 0.25μm).The temperature of the programme was 40°C isothermal time; heating up to 250°C with a heating rate of 40°C/min. Helium was used as carrier gas with a flow rate of 1.0 ml /min. 1μl sample insertion volume was employed. The inlet temperature was controlled as 250^°^C. The MS source temperature was 230°C.

### 3.6. Characterization of ZnO nanoparticles

The synthesized nanoparticles were characterized by various techniques. They are SEM, XRD analysis, FTIR spectroscopy and UV–Vis spectroscopy.

## 4. Result And Discussion

The results of morphological characteristics of *Rhizobacteria*l isolates were summarized in **Table 1** The *Rhizobacteria*l isolates were appear dominant growth on YEMA medium and two isolates were designated as RN00A and RN00B. The isolates were slow growers and growth obtained after 3-4 days. The RN00A failed to absorb Congo red in the medium.

**Table 1.** shows the morphological characterization of Rhizobacteria

| Isolate name | Colony characteristics on YEMA |  |  |  |  |  |  |
| --- | --- | --- | --- | --- | --- | --- | --- |
|  | Size | Shape | Texture | Opacity | Pigmentation | Margin | Elevation |
| RN00A | Medium | Circular | Mucoid | Translucent | White | Entire | Raised |
| RN00B | Small | Circular | Smooth | Opaque | Pink | Undulate | Flat |

The RN00A colony morphology on YEMA medium was mostly circular, mucoid, white and translucent. They showed highly mucoid on after 5 days of incubation at 26°C. The RN00B colony was circular, smooth, pink, and opaque.

### Biochemical characterization of Rhizobacteria

The biochemical characteristics of the isolated organisms are summarized in the **Table 2**. The bacterial isolate RN00A was aerobic, Gram-negative, rod-shaped, and motile. The isolate RN00A showed positive reactions in Methyl Red (MR), Starch hydrolysis, Triple Sugar Iron (TSI) agar, Nitrate reduction, Glucose Peptone Agar (GPA), Catalase, Oxidase, Urease test and Lactose assay. They were negative to Indole production, VogesProskauer (VP), Citrate utilization, Gelatin hydrolysis, Keto - lactose test, and Flourescent assay. The bacterial isolate RN00B was aerobic, Gram-positive, rod- shaped and motile. The cells contained endospores. The isolate RN00B showed positive reactions in Indole production, Citrate utilization, Starch hydrolysis, Triple Sugar Iron (TSI) agar, Gelatin hydrolysis, Glucose Peptone Agar (GPA), Oxidase, Catalase test and Lactose assay. They are negative to Methyl Red (MR), Voges - Proskauer (VP), Urease, Nitrate reduction, Keto - lactose test, and fluorescent assay.

**Table 2.**
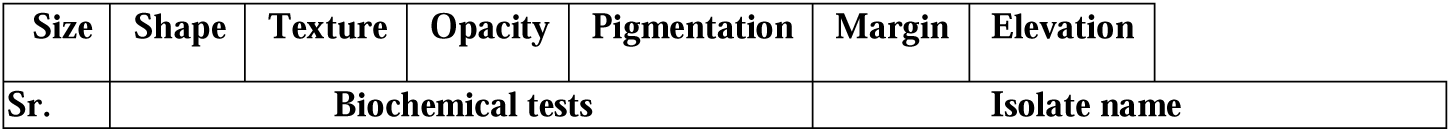
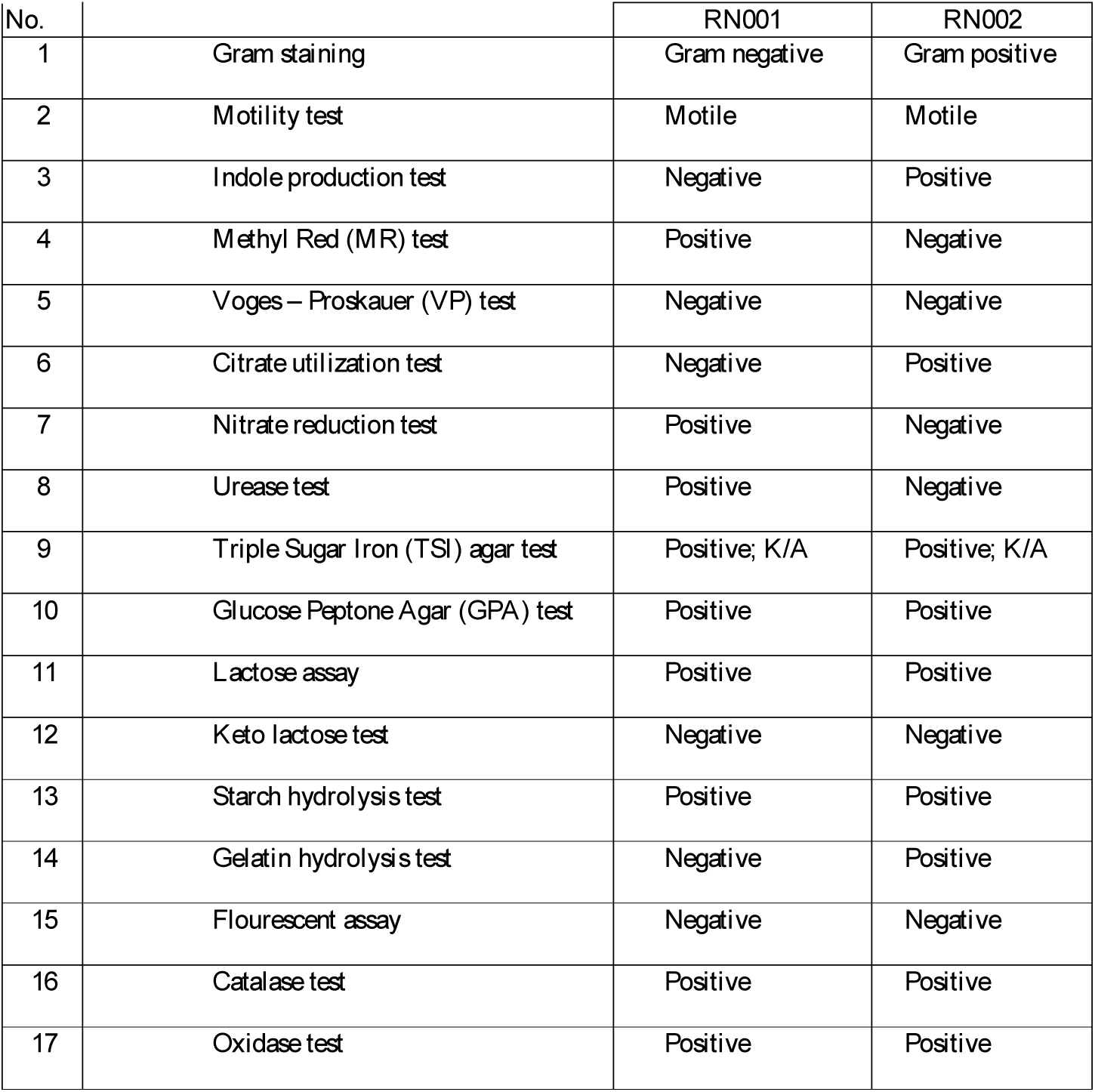
Biochemical characterization of Rhizobacteria

### Antibacterial Activity

DMSO was served as negative control and standard antibiotic solutions like Amoxicillin (AX) and streptomycin (SM) was used as positive control. The antibacterial activity of different plant extracts of Vigna Mungo in different test organisms showed antibacterial activity in the range between 14 to 24mm.The methanol extract (VMNP) showed highest zone of growth inhibition against B. zanthoxyli (24 mm) followed by B. velezensis (14 mm). The antibacterial activity of green synthesized zinc oxide nanoparticles in different test organisms showed broad spectrum antibacterial activity in the range between 13 to 25 mm. Among them (VMNP) showed highest zone of growth inhibition against K. ozaenae (25 mm), B. zanthoxyli (21 mm) and B. velezensis (15 mm).The antibacterial activity of Rhizobacterial synthesized NPs showed antibacterial activity in the range between 11 to 22 mm against test organisms. Among them the pellet of both Rhizobium sp. (NWP) and Bacillus flexus (NPP) showed broad spectrum antibacterial activity in the range between 15 to 22 mm. Microbial synthesized zinc oxide nanoparticles showed greater antimicrobial potential when compared with chemically synthesized zinc oxide nanoparticles..

### Minimum inhibitory concentration (MIC)

The MIC was determined against two bacterial strains such as *B. velezensis* and *K. ozaenae*. The NPS synthesized from the pellet of *Rhizobium* sp. expressed minimal MIC against *B. velezensis* (0.04 µg/mL) followed by *K. ozaenae* were showed in Table 3.

**Table 3.** MIC results

| Concentration<br>(µg/mL) | Diameter of zone of inhibition (in mm) |  |
| --- | --- | --- |
|  | *B. velezensis* | *K. ozaenae* |
| 0.02 | nil | nil |
| 0.04 | 12 | 15 |
| 0.06 | 19 | 20 |
| 0.08 | 23 | 22 |

### Minimum bactericidal concentration (MBC)

The MBC results revealed that zinc oxide nanoparticles of *Rhizobium* sp. significantly inhibit the growth of selected test organisms (table 4). The zinc oxide nanoparticles expressed minimal MBC value against *Bacillus velezensis* (0.5 µg/mL) followed by *Klebsiella ozaenae*.

**Table 4.** MBC results

| Concentration<br>(µg/mL) | Number of colonies |  |
| --- | --- | --- |
|  | *Bacillus velezensis* | *Klebsiella ozaenae* |
| Control | TNTC | TNTC |
| 0.06 | 326 | TNTC |
| 0.07 | 262 | 298 |
| 0.08 | 125 | 280 |
| 0.09 | 63 | 119 |
| 0.1 | 29 | 60 |
| 0.5 | nil | nil |

### GC-MS Analysis

The spectrum of the compounds was correlated with the database of spectrum of identified compounds were stored in the GC-MS library. The summary of analysis is given in Table 5. The GC-MS study of leaf extract of *Vigna Mungo* plant have shown many phytochemicals. The major components present in the methanol extract of *V. mungo* along with the molecular formula, molecular weight, and retention time are presented in Table 5. The methanol extract of *Vigna Mungo* leaves showed the content of Phytol, 2,6,10,15,19,23 –hexamethyl tetracosane and Alpha- tocopherol. These compounds have been frequently attributed to their antioxidant, antitumor and antimicrobial activity ^[7]^. Vigna have antioxidant properties and can manage and cure different diseases linked with free radical generation _[9]_.

**Table 5.**
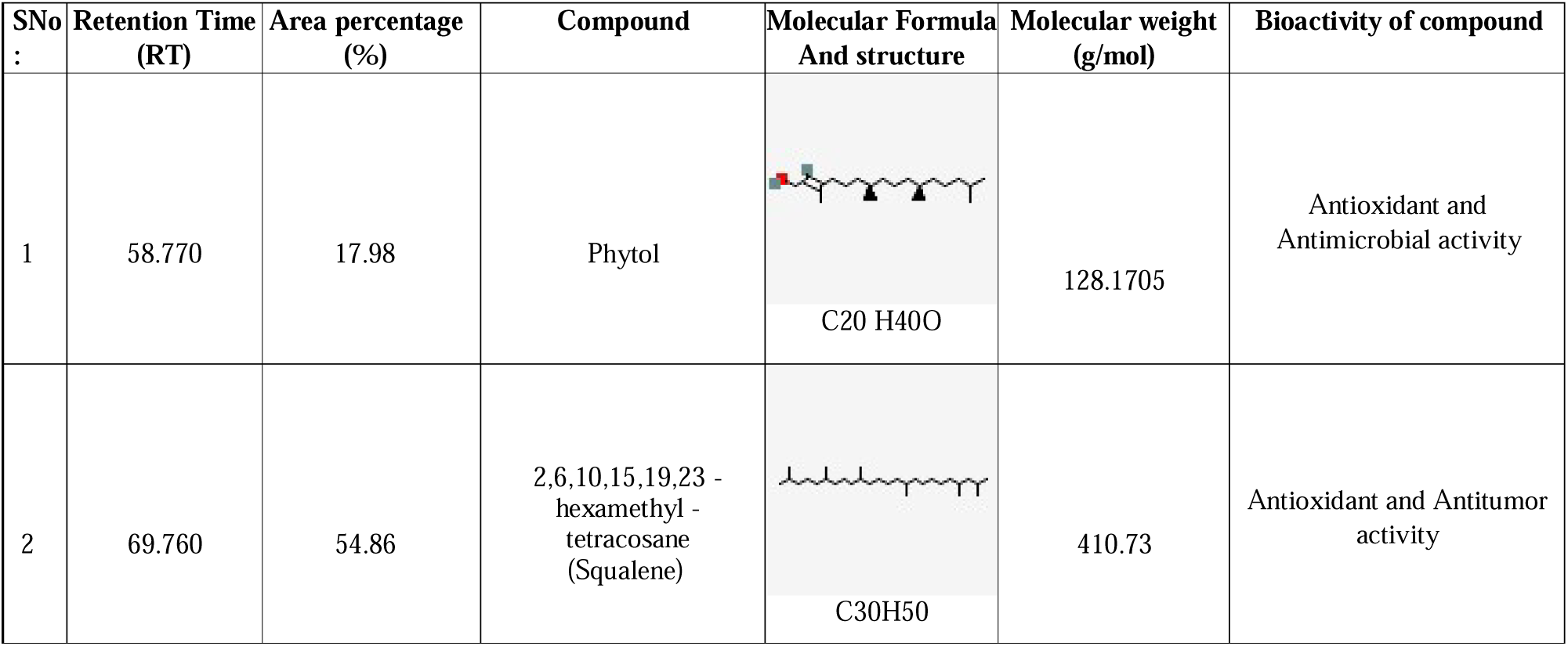
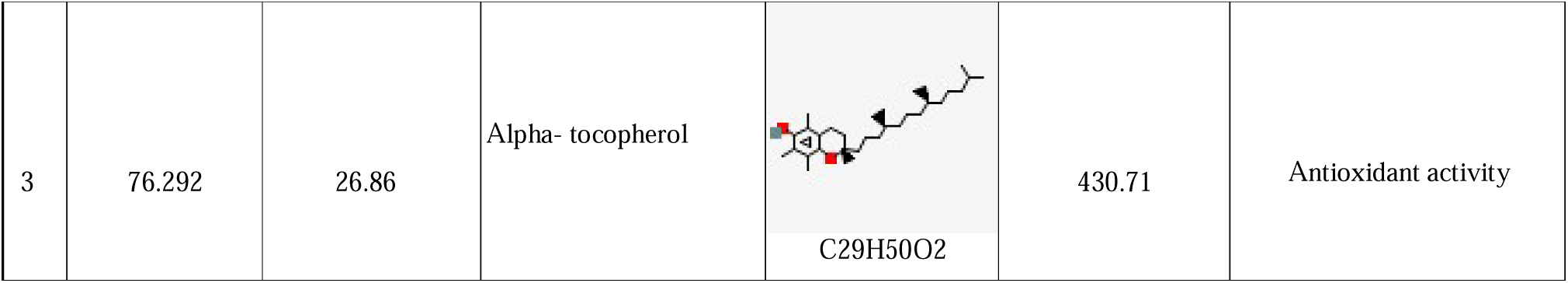
GC-MS Analysis of Vigna Mungo leaf extract

| SNo<br>: | Retention Time<br>(RT) | Area percentage<br>(%) | Compound | Molecular Formula<br>And structure | Molecular weight<br>(g/mol) | Bioactivity of compound |
| --- | --- | --- | --- | --- | --- | --- |
| 1 | 58.770 | 17.98 | Phytol | C20 H40O | 128.1705 | Antioxidant and<br>Antimicrobial activity |
| 2 | 69.760 | 54.86 | 2,6,10,15,19,23 -<br>hexamethyl -<br>tetracosane<br>(Squalene) | C30H50 | 410.73 | Antioxidant and Antitumor<br>activity |
| 3 | 76.292 | 26.86 | Alpha- tocopherol | C<sub>29</sub>H<sub>50</sub>O<sub>2</sub> | 430.71 | Antioxidant activity |

### Characterization of ZnO nanoparticles

These images demonstrated that zinc oxide nanoparticles are spherical in shape and each particle was aggregation of many smaller particles ^[3]^. Claimed that the morphology of ZnO nano-powder using zinc acetate is smoother than in zinc nitrate. Moreover, precursor concentration plays a great role on morphological features of nanoparticles. According to Pourrahimi *et al*. (2014), ZnO can be form in different structures due to the type of precursors that has been used ^[11]^.

### XRD analysis

The XRD pattern of ZnO NPs synthesized by the *Rhizobium* sp. XRD shows 2θ values at 31.548°, 34.197°, 36.017°, 47.292°, 56.360°, 62.622°, 66.15°, 67.729°, 68.820°, 72.3090 and 76.7390. It also confirms the synthesized nano powder was free of impurities as it does not contain any characteristics XRD peaks other than zinc oxide peaks.

### FTIR spectroscopy

FTIR spectra (of the synthesized nanoparticles were recorded in the 500–4500 cm^_1^ range. The absorption at 1227 cm^-1^ indicates the presence of hydroxyl group. The absorption at 1687 indicates the presence of alkane groups. The absorption at 1593 indicates the presence of carbonyl groups. The weak absorption at 2015 is due to C=C stretching vibration. The absorption peaks at 2686 and 3120 indicates the presence of amide and aromatic alcohol. They could possibly enhance the stabilization of ZnO nanoparticles in the aqueous medium.

### UV-VIS spectroscopy

Graph 3 shows the UV-Vis absorption spectrum of zinc oxide nanoparticles. The absorption spectrum was recorded for the sample in the range of 280 - 420 nm. The spectrum showed the absorbance peak at 288 nm corresponding to the characteristic band of zinc oxide nanoparticles. On the surface of nanoparticles, the electron clouds are present which are able to oscillate and absorbs the electromagnetic radiation at a particular energy, energy corresponding to the photons of 288 nm. This resonance is known as surface plasmon resonance (SPR).

## 5. CONCLUSION

The current work demonstrates that zinc oxide nanoparticles were effectively generated using a biological technique. Rhizobacteria may be used as a biological system to readily produce ZnO NPs. Rhizobium sp., for example, may be regulated under controlled circumstances and has a high potential for extracellular and intracellular metallic nanoparticle formation. This approach is straightforward, inexpensive, and free of pollutants and pollution, producing a huge volume of stable nanoparticles. According to the findings of the preceding investigation, zinc oxide has antibacterial properties. Nanoparticles and V. mungo leaf extract are also confirmed. Bacillus velezensis, Bacillus zanthoxyli, and Klebsiella ozaenae were used as test microorganisms for antibacterial activity. The presence of Phytol, Squalene, and Alpha - tocopherol is revealed by GC-MS analysis of methanol leaf extract of V. mungo. These findings show that ZnO nanoparticles and V. mungo leaf extract have a broad spectrum of antibacterial properties against a variety of possible harmful bacteria and might be used in nanodrug formulations. On the basis of the general findings of this investigation, We may infer that biosynthesis of ZnO NPs using V. mungo and Rhizobacteria is significantly safer and more environmentally friendly than physical and chemical approaches.

## 6. ETHICS APPROVAL AND CONSENT TO PARTICIPATE

Not applicable.

## 7. HUMAN AND ANIMAL RIGHTS

No Animals/Humans were used for studies that are base of this research.

## 8. CONSENT FOR PUBLICATION

Not applicable.

## 9. AVAILABILITY OF DATA AND MATERIALS

The author confirms that the data supporting the findings of this research are available within the article.

## 10. FUNDING ACKNOWLEDGEMENT AND CONFLICT OF INTEREST

The authors whose names are listed immediately above certify that they have NO affiliations with or involvement in any organization or entity with any financial interest (such as honoraria; educational grants; participation in speakers’ bureaus; membership, employment, consultancies, stock ownership, or other equity interest; and expert testimony or patent-licensing arrangements), or non-financial interest (such as personal or professional relationships, affiliations, knowledge or beliefs) in the subject matter or materials discussed in this manuscript.

